# Exploratory and risk-taking behaviours in coexisting rodents

**DOI:** 10.1101/2024.01.09.574853

**Authors:** Bryan Hughes, Jeff Bowman, Albrecht Schulte-Hostedde

## Abstract

There has been an increasing interest in modelling the influence of animal personality on species interactions within ecosystems. Animal personality traits associated with dispersal, movement within a home range and risk-taking, including docility and exploration, have been shown to influence an array of environmental variables including seed dispersal and habitat availability. Despite growing interest however, little information is available to model the effects of differences in personality phenotypes among coexisting species. Since coexisting or sympatric species often compete for resources, differences in movement patterns can help mitigate the impact of intra- and interspecific competition. We used two standardized behavioural tests with three species of coexisting rodents in Algonquin Provincial Park, Ontario, Canada to measure exploration and docility personality phenotypes. To evaluate personalities, we modelled plastic changes in behaviours within species and phenotypic variation in behavioural strategies among species. We show empirical evidence to support differences in personality phenotypes in coexisting species and consider the importance of alternative personality strategies in shaping community dynamics.

## Introduction

Genetic and phenotypic variation in the life-history, behaviour and physiology of a species are mediated through natural selection and ecological trade-offs in resource acquisition and allocation (Roff and Fairbairn, 2007). The pace-of-life syndrome hypothesis (POLS) is an analytical framework that suggests the life history, behaviour and physiology of an organism exhibits predictable correlations between traits (Reale et al., 2010; Dammhahn et al., 2018).

Thus, the patterns suggested by the POLS hypothesis follow a fast-slow continuum, where short-lived organisms with a high reproductive output demonstrate proactive or “fast” behavioural and physiological phenotypes. In contrast, long-lived organisms that have fewer offspring but invest more in the care of individual young demonstrate more reactive or “slow” phenotypes (Réale et al., 2010). While there is empirical evidence to support differences among species along the fast-slow continuum (Ricklefs and Wikelski, 2002), evidence to support the POLS hypothesis among individuals of the same species remains mixed (Dammhahn et al., 2018). For example, the relationship between boldness and reproductive success, alongside the costs of survival has been shown, however specific mechanisms evaluating personality and trade-offs remain unclear (Smith and Blumstein, 2008; Biro and Stamps, 2008). One reason for the lack of information on specific relationships within populations results from the influence of environmental factors (Bielby et al., 2007; Dammhahn et al., 2018; Royaute et al., 2018; Hamalainen et al., 2021). By further investigating differences in animal personality and behaviour, we may gain additional insight into how species interact within an ecosystem.

An extensive body of literature on the personality of various rodent species shows plastic changes in traits associated with the POLS between individuals and variation among species (Dingemanse et al., 2010; Brehm et al., 2020). Within an ecosystem, trade-offs in exploration and risk avoidance have been shown to promote species coexistence (Morris and Palmer, 2023; Milles et al., 2020). Such intraspecific differences in personality, specifically traits related to movement, have been shown to regulate community dynamics (Milles et al., 2020).

Relationships between POLS traits are heavily influenced by trade-offs associated with individual reproduction (Healy et al., 2019). Reproduction is an energetically expensive and potentially risky investment, and rodents may invest in seasonal or continuous breeding strategies to accommodate energetic costs (Bergeron et al., 2011; Bronson and Perrigo, 1987). Although many rodents may breed year-round if environmental conditions are favourable, seasonal breeding strategies are common in populations that experience consistent periods of resource scarcity – such as winter (Wolff and Sherman, 2008). Such Seasonal breeding events have been linked to plastic changes in personality (Le Coeur et al., 2015; Eccard and Herde, 2013). For seasonally breeding rodents, both males and females experience an increase in hormone levels that facilitate mate acquisition and selection during the breeding season (Kavaliers and Choleris, 2017). Given the associations predicted by the POLS hypothesis, variation in physiology during active breeding periods should result in a shift in behaviours associated with mate acquisition.

Deer mice (*Peromyscus maniculatus*), red-backed voles (*Clethrionomys gapperi*), and woodland jumping mice (*Napaeozapus insignis*) are three rodent species that often inhabit a similar ecological niche (Fryxell et al., 1998). Each of these three species expresses a polyandrous mating system and have an active breeding season from May through August during which individuals may breed several times. Despite similarities, each species has varying life history traits that may be modelled along the fast-slow continuum outlined by the POLS hypothesis. Deer mice generally display fast behaviours (Careau et al., 2011), have litter sizes averaging 4-6 pups (Maser et al., 1981), and have an average lifespan of 1-2 years (Dice, 1936). Woodland jumping mice live longer than voles and deer mice, living up to 4-5 years (Wrigley, 1972). Woodland jumping mice also have a lower reproductive output than either deer mice or red-backed voles, engaging in only 1-2 breeding events per season. While jumping mice have a greater gestation and weening period than both deer mice and red-backed voles, Jumping Mice only produce 2-4 pups per litter (Whitaker and Wrigley, 1972). Red-backed voles live 1-2 years and produce litter sizes that range from 4-5 pups, up to 2-3 times per reproductive season (Merrit, 1981). Based on the lifespan and reproductive output of these three species, deer mice should express the “fastest” behaviours, while jumping mice should express “slower” behaviours, meanwhile red-backed voles may express behavioural traits that are intermediate compared to deer mice and woodland jumping mice.

Because animal personality is a driving mechanism for determining how individuals and species react to environmental influences (Réale et al., 2010), it is important to understand how different behavioural strategies arise among species during different parts of the reproductive cycle. A home range refers to the spatial area where an individual may forage for resources or compete for mate selection. Dispersal and movement patterns are the predominant mechanisms to mitigate potential intraspecific competition in mate selection; therefore, reproductive output is heavily influenced by dispersal and movement patterns within a species home range (Perrin and Maxalov, 2000). Evaluating behaviours that are associated with movement during different stages of a species reproductive season may help provide insight into factors mediating the expression of different behavioural strategies. In rodents, docility is considered a measure of how an individual reacts to a potential predatory or risky situation (Boice et al., 1968; Martin and Réale, 2008), whereas exploration is a measure of the engagement or movement of an individual within the environment (Careau et al., 2009). Therefore, actions that reflect docility and exploration are both indicators of an individual’s movement potential and likelihood to engage with novel components of an environment (Dingemanse et al., 2003; Réale et al., 2007; Morris 1984; Petelle et al., 2013).

Despite the relevance of personality as a driving mechanism for species interactions between conspecifics, other species and the local environment, there are few and often inconclusive studies evaluating the expression of personality in wild populations (Réale and Dingemanse, 2012). Our study aims to evaluate behavioural strategies among species during different stages of the reproductive cycle. We focused on three species: the deer mouse, the red-backed vole, and the woodland jumping mouse. Similarly, we evaluated intraspecific differences in movement behaviour throughout the breeding season. Because each of these three species should benefit from the same correlation of traits, such that less docile individuals explore more, we predicted that behavioural syndromes that influence habitat use should arise such that more explorative species and individuals should also express less docile personality traits. Despite variation in microhabitat usage (Schulte-Hostedde and Brooks, 1997) which also reduces intra-specific competition, populations of these three species often compete for similar habitat and resources when populations occupy the same area (Davidson and Morris, 2001; Dracup et al., 2016). We hypothesized that if coexistence promotes alternative behavioural strategies to accommodate intra- and interspecific competition, then there should be measurable differences in personality among species along the proposed “fast” to “slow” continuum. Indeed, Distinct differences in personality have been shown to arise as a co-adaptive set of behaviours among species, potentially enabling coexistence (Morris and Palmer, 2023). We further predicted that there should be measurable variation in behaviours relating to movement patterns among species. Therefore, observable differences in behavioural strategies among species and between individuals should arise during the active breeding season, such that some species or individuals will be more explorative and less docile. Further among different species, we predicted similar directionality of syndromes at different positions along the predicted “fast -slow” continuum.

Among individuals of the same species, we predicted that actively breeding males should express a higher rate of proactive behaviours compared to non-breeding males. Likewise, females that are actively investing in the care or development of young should express more reactive behavioural strategies, given the energetic investment of maternal care (Rémy et al., 2011).

## Methods

### Sampling

Deer mice, woodland jumping mice and red-backed voles were collected from May-September 2022 under animal handling procedures approved by Laurentian University (protocol #6011106). All animals were sampled from Algonquin Provincial Park, Ontario, Canada (45°54′ N, 78°26′ W) from 17 established traplines (Fryxell 1998). Sherman traps (H.B. Sherman Traps, Inc., Tallahassee, Florida) were set bi-weekly at dusk with water-soaked sunflower seeds over 3 consecutive nights and were checked at dawn. Because trap confinement may have an impact on individual behaviour (Brehm et al., 2020), each trapline was checked in the same order each morning, and the start time of any behavioural test was recorded to estimate the impact longer confinement may have on behaviour. To measure the potential impact of cumulative testing and handling on behaviour, we maintained a record of the total number of times any individual was captured (denoted as cumulative capture) and the total number of times an individual entered a behavioral assay throughout the season. Captured individuals were collected from the Sherman traps in a plastic handling bag. Individuals were weighed with a Pesola scale (+/- 0.1 g), sexed, assigned an age class (juvenile, sub-adult, adult) based on mass and identifiable morphological characteristics (Schmidt et al., 2019). Breeding status was assessed in individuals as scrotal or non-scrotal for males, and pregnant, lactating, or non-reproductive for females. All behavioural testing and animal handling took place directly in the field after capture.

### Behavioural assays

Captured individuals were subject to either a 1-minute Handling Bag Test or a 5-minute Open Field Test each trap day. Handling Bag Tests are used to measure behaviours associated with animal docility (Martin and Reale 2008), while Open Field Tests are used to measure behaviours associated with exploration and activity (Carter et al., 2013). All behavioural assays were recorded in the field using a video camera (Sony HDR-CX405), and behaviours were later analyzed using the recording. The Handling Bag Test was conducted before animals were weighed and measured. Individuals were left in the plastic handling bag for one minute and held at arm’s length from the observer. Docility during the Bag Test was measured as the total amount of time an individual spent motionless (Martin and Réale, 2008; Supplementary data A).

The Open Field Test was conducted inside a novel plastic arena (51 x 41 x 74 cm) fitted with an 8.89 cm PVC opening and a mesh barrier on top to prevent an individual from leaving the arena, while still allowing for recording. Arena tests were filmed for a total of five minutes starting from when the individual entered the arena through the PVC pipe opening. Total exploration behaviour was measured as the time an individual spent moving around the arena or engaging with the arena through scratching or sniffing at the plastic, while non-exploratory time was measured as time spent motionless either in the arena, or the entrance marked as hiding behaviour (Supplementary data B). The arena was cleaned between each trial using an 80 % vinegar solution and then rinsed with water. For either the Handling Bag Test or the Open Field Test, grooming was considered a separate behaviour and not counted towards the total of either docile or explorative personality type.

### Video processing

All videos were recorded and later assessed using CowLog 3.0 (Pastell 2016) to quantify behaviours using a pre-defined ethogram (Supplementary data A and B). We analyzed 222 Handling Bag Tests and 156 Open Field Tests.

### Statistical analyses

Statistical analyses were conducted using statistical software analyses packages in R, version 4.2.3 (R Core Team 2023). We ran a repeatability analysis on behavioural variables for each species to determine if the behaviours observed in each video were repeatable across individuals and thus considered personality phenotypes (Wilson 2018). Because there were no repeated observations of Jumping Mice with the exploration test, we were unable to test for repeatability for this test. Each repeatability test was performed using the rptR package in R (Stoffel et al., 2017), using reproductive condition, age and sex as fixed effects, and individual ID as a random effect. Because recapture events may increase acclimation to behavioural tests, individuals that were retagged due to lost or unidentifiable tag numbers were removed from analyses, as cumulative capture count would be uncertain. We did not limit the repeatability test to individuals with only one observation (single capture count) because this has been shown to miss variation in individual plasticity (Martin 2011).

To evaluate the relationship between exploration and docility, we ran Spearman’s rank correlation tests between the first Bag Test and Open Field Test for each individual. Because this relationship is dependent on the same individual undergoing both the Bag Test and Open Field Test, only individuals that had undergone both tests within the same week were used for this analysis (n = 40 deer mice, 16 red-backed voles, and 4 woodland jumping mice). We only used the first observation of each test to reduce the impact of acclimation or learned behaviour from repeated testing (Cnops et al., 2022). To reduce the impact of temporal variables, we only used individuals that underwent both behavioural tests during the same 3-day trapping period.

Because retagged individuals may have already been subject to behavioural testing, individuals marked as retagged or having missing ear tag numbers were also excluded from this section of the analysis.

To measure the impact from handling and morphometric variables on personality, we used a generalized linear mixed effects model using the package MCMCglmm (Hadfield, 2010) to test our predictions on exploration and docility personality phenotypes. For each species separately, we used personality as our response variable, with either age, sex, reproductive status, body mass, or capture count, date and start time as fixed effects, with individual ID as a random effect. For the exploration personality trait, we used ‘tagged’ as an additional variable to represent if an individual received an ear-tag prior to entering the Open Field Test. Since only adult individuals were recorded for woodland jumping mice, we did not include age as a fixed effect for those models. Akaike information criterion (AIC) corrected for small sample sizes (AICc), and likelihood-based pseudo-R-squared (LR Χ^2^) were calculated using the package MuMIl version 1.47.5 (Barton, 2023).

We then used a linear mixed effect model from the R package lmer4 (Bates et al., 2015) for each species and behavioural test pooled together using each personality test as a dependent variable. We used reproductive condition, age, weight, capture count and date as a fixed effect, and individual ID as a random effect. For this model, sex and reproductive condition were not included together because the two categories are dependent.

## Results

### Repeatability estimates

We analyzed 202 Bag Tests, including observations of 83 deer mice (135 total tests), 48 red-backed voles (65 total tests) and 15 woodland jumping mice (16 total tests). We also analyzed 147 Open Field Tests including observations of 84 deer mice (93 total tests), 30 red-backed voles (36 total tests) and 14 woodland jumping mice (18 total tests). For the Bag Test, docile behaviours were repeatable in deer mice, but not in red-backed voles or woodland jumping mice. For the Open Field Tests, non-exploration behaviour was repeatable in deer mice and red-backed voles (Table 1).

**Table 1:** The results from the repeatability analysis for the (Bag Test) and (Open Field Test) measuring Docility and Exploration respectively. Results highlighted in bold were determined to be repeatable. All estimates were calculated using mixed effects model with age, sex, and reproductive condition as a fixed effect and individual ID as a random effect. 95% confidence intervals were calculated using parametric bootstrapping.

| Docility - ID random effect |  |  |  |  |  |
| --- | --- | --- | --- | --- | --- |
| Behavioural Variable | Mean | R | SE | CI (95%) | P |
| deer mouse |  |  |  |  |  |
| **Movement** | **0.341** | **0.269** | **0.143** | **0.0351, 0.6** | **0.044** |
| Escape | 0.148 | 0.066 | 0.13 | 0, 0.443 | 0.33 |
| Grooming | 0.0998 | 0.031 | 0.093 | 0, 0.313 | 0.35 |
| Foraging | 0.109 | 0 | 0.122 | 0, 0.407 | 1 |
| **Freezing** | **0.404** | **0.339** | **0.137** | **0.116, 0.655** | **0.020** |
| red-backed vole |  |  |  |  |  |
| Movement | 0.34 | 0 | 0.268 | 0, 0.837 | 1 |
| Escape | 0.369 | 0 | 0.259 | 0, 0.805 | 1 |
| Grooming | 0.348 | 0 | 0.243 | 0, 0.761 | 0.5 |
| Freezing | 0.354 | 0 | 0.268 | 0, 0.819 | 1 |
| woodland jumping mouse |  |  |  |  |  |
| Movement | 0.0001 | 0 | 0 | 0, 0 | 1 |
| Escape | 1 | 1 | 0 | 1, 1 | 0.30 |
| Freezing | 1 | 1 | 0 | 1, 1 | 0.30 |

Exploration - ID random effect
| Behavioural Variable | Mean | R | SE | CI (95%) | P |
| --- | --- | --- | --- | --- | --- |
| deer mouse |  |  |  |  |  |
| Move | 0.284 | 0.066 | 0.207 | 0, 0.691 | 0.37 |
| Groom | 0.237 | 0 | 0.214 | 0, 0.66 | 1 |
| **Freezing** | **0.842** | **0.809** | **0.067** | **0.695, 0.943** | **0.0019** |
| Hide | 0.227 | 0 | 0.209 | 0, 0.658 | 1 |
| red-backed vole |  |  |  |  |  |
| Move | 0.836 | 0.315 | 0.232 | 0, 1 | 0.34 |
| Groom | 0.741 | 0 | 0.308 | 0, 1 | 0.5 |
| Freezing | 0.856 | 0.391 | 0.205 | 0.187, 1 | 0.097 |
| **Hide** | **1** | **1** | **0** | **1, 1** | **0.000067** |

### Correlation between docility and exploration

Our results suggested that the relationship between docility and exploration in deer mice (n = 40) and red-backed voles (n=16) was non-significant. We report a non-significant negative relationship between docility and exploration in red-backed voles (r = -0.437, P = 0.091), and a lack of any relationship between these personality types in deer mice (r = -0.018, P = 0.26).

### Docility and exploration models

For models predicting docility among species, age, sex and reproductive status were only substantially better than the null model in deer mice. For exploration models, age, sex, reproductive status and handling variables capture count, date, start time and tagged were stronger than the null model (Table 2). However, there was a negative correlation (Rs = -0.28, P = 0.0066) between docile behaviour and cumulative capture count in deer mice. Non-docile behaviour also had a significant relationship with reproductive condition in deer mice, but neither woodland jumping mice or red-backed voles.

**Table 2:**
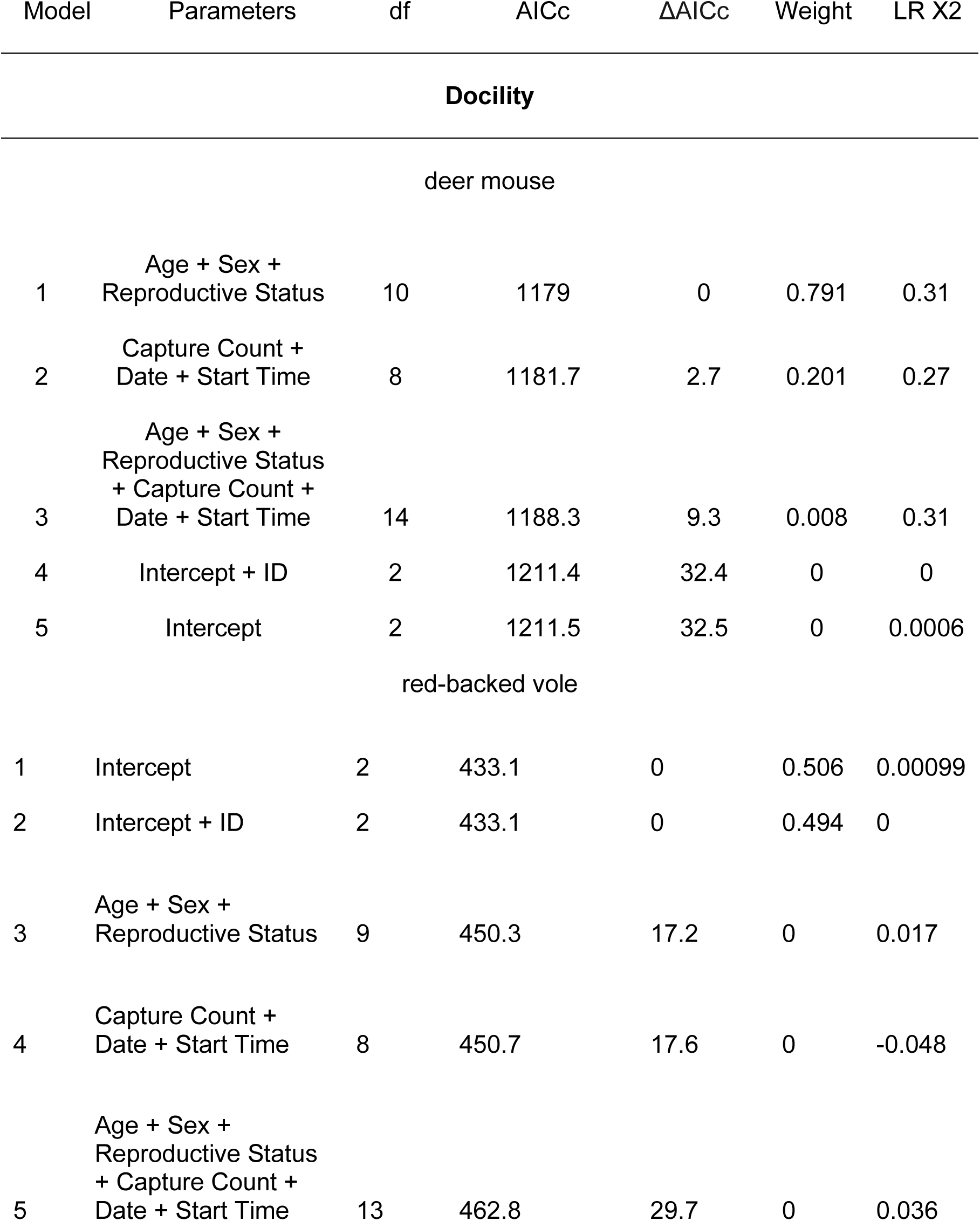
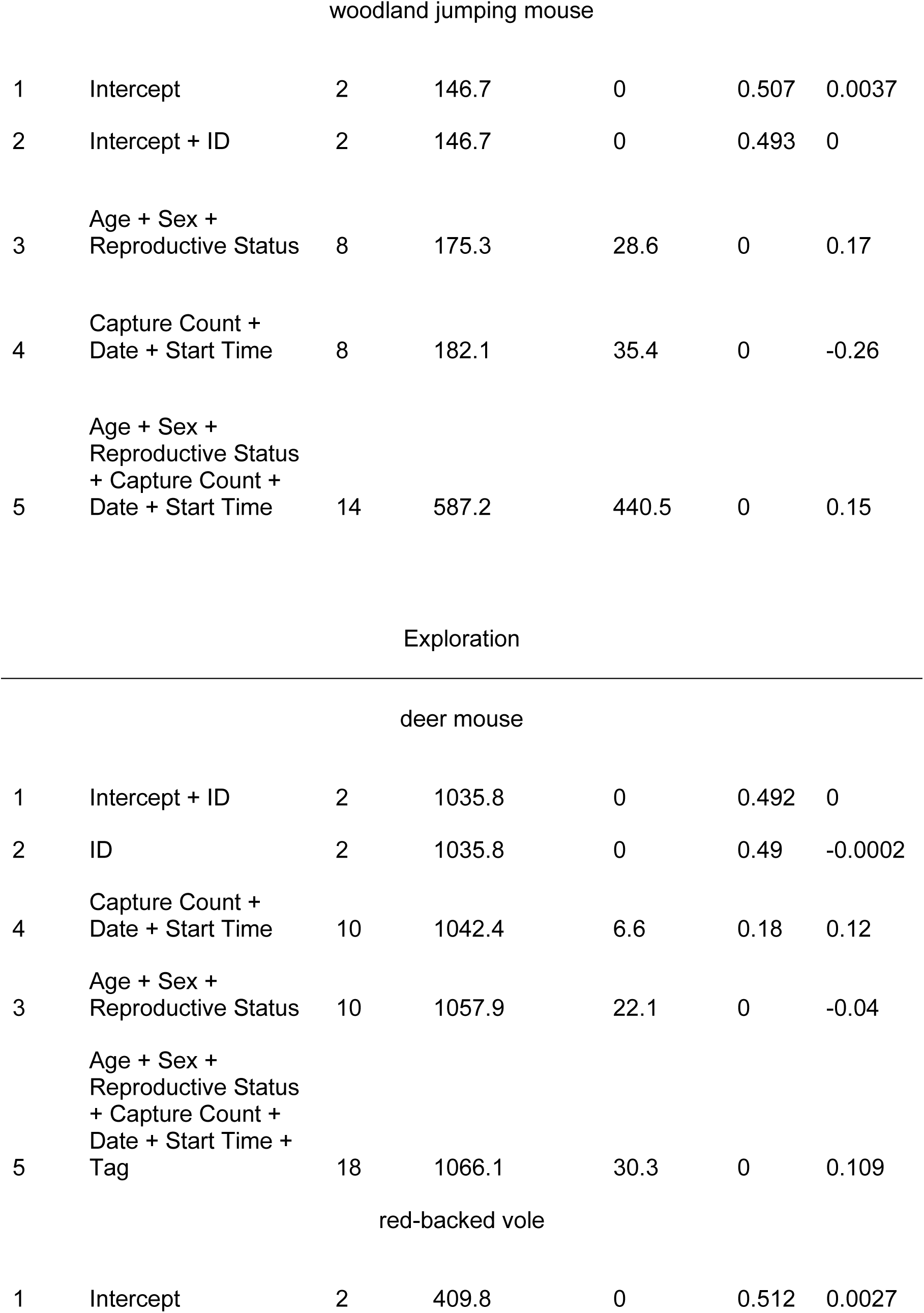
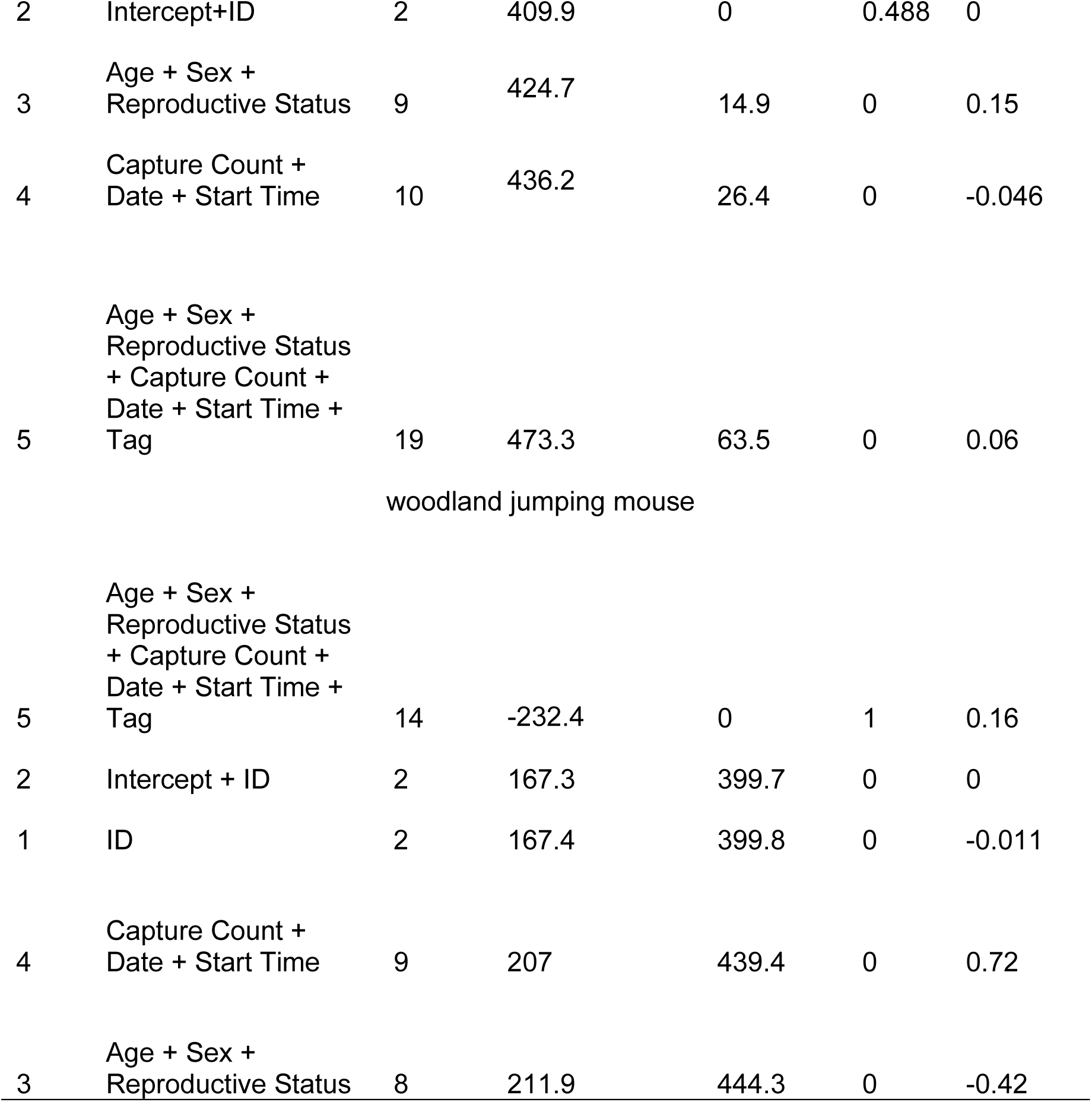
Summary of bivariate Bayesian models examining the relationship between personality type (docility or exploration) within each species (docility, deer mouse n = 135, red-backed vole n = 65, woodland jumping mouse = 16; exploration, deer mouse n = 93, red-backed vole n = 35, woodland jumping mouse n = 14) using individual ID as a random effect in all models.

**Table 3:**
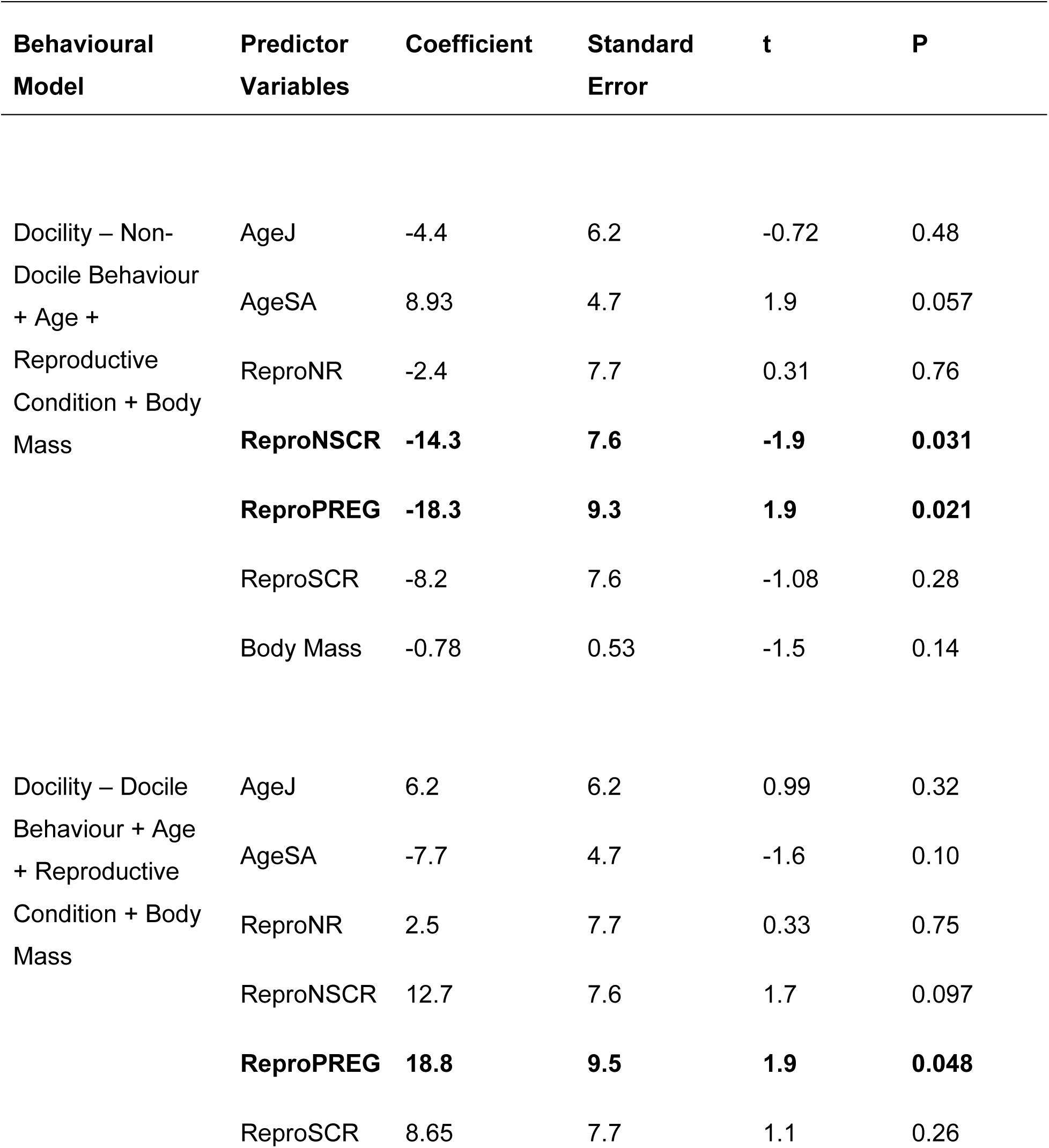
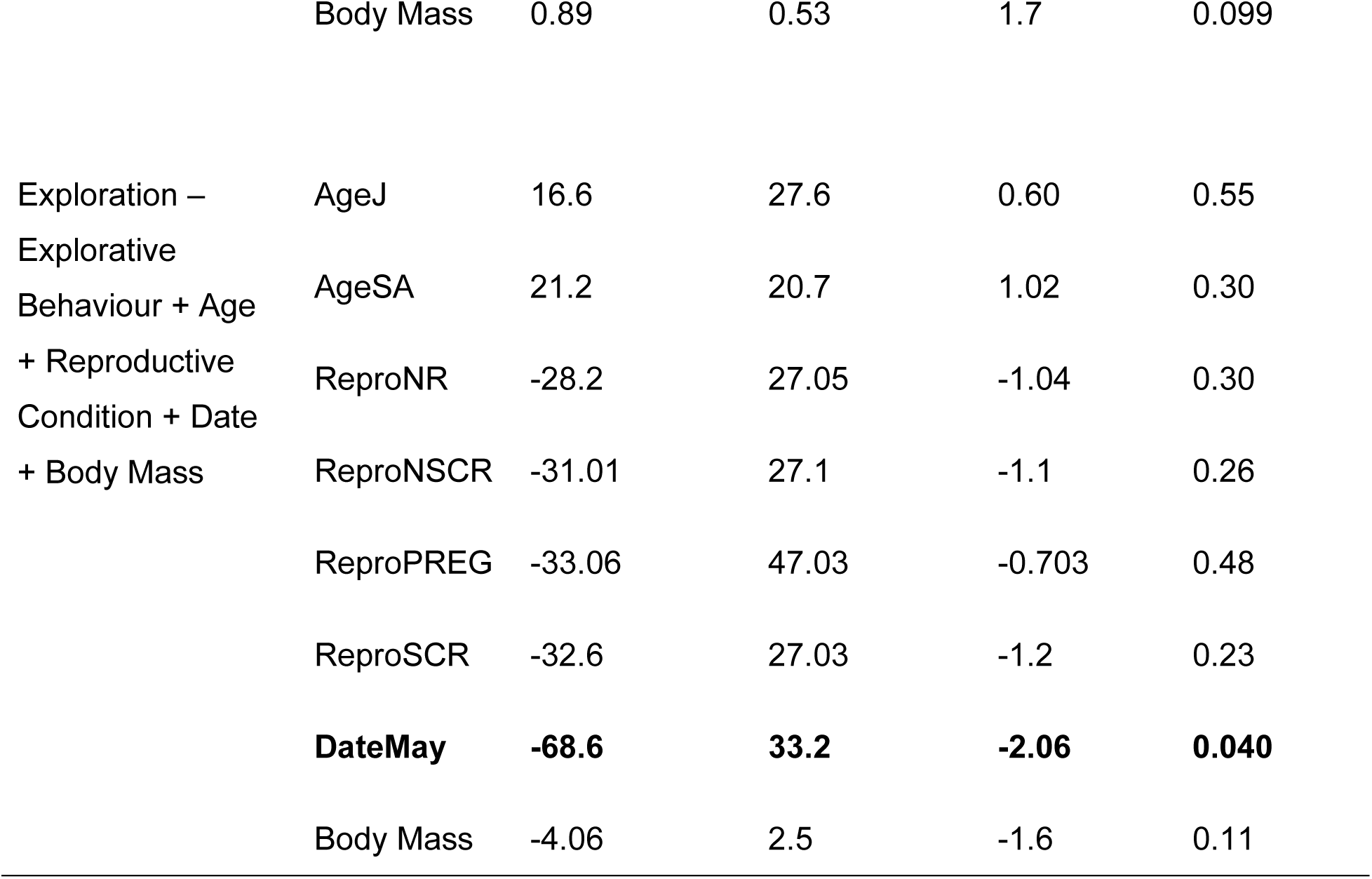
Summary of top best fit linear mixed effects models predicting the relationships between personality (docility, measured the amount of time an individual spends trying to escape versus remaining motionless; or exploration measured as the amount of time an individual moves around a novel environment versus remaining motionless or hidden) pooled across species (n = 202 Bag Tests and 147 Open Field Tests).

### Cross-species comparisons of docility and exploration

The linear effects model showed significant differences in docility among species during different stages of the reproductive cycle (F = 4.8, df = 216, P = < 0.001, R^2^ = 0.081). Non-scrotal males (P = 0.031) and pregnant females (P = 0.021) had a significant relationship with explorative behaviours. Docile behaviour also had a significant relationship with reproductive condition (F = 11.67, df = 214, P = 0.026, R^2^ = 0.25), such that pregnant (P = 0.048) individuals spent more time remaining motionless, thus expressing more docile behaviour (Table 4). Despite high variation between individuals, there were distinct differences among species during each reproductive phase. Deer mice consistently expressed the lowest amount of docile behaviour, while red-backed voles displayed the greatest amount of docile behaviour. Woodland jumping mice consistently expressed an intermediate amount of non-docile behaviour compared to deer mice and red-backed voles across all reproductive phases (Figures 2 and 3). There were no significant relationships between exploratory behaviour (time spent moving) in the Open Field Test and any of the predictor variables - sex, age, reproductive condition, month of capture, date of receiving an ear tag - when using individual ID as a random effect (F = 2.87, df = 141, P = 0.0014, R^2^ = 0.19) Using species as an independent variable, there was a positive correlation between capture count (r = 0.22, P = 0.010) and exploration (n = 93 deer mice, 44 red-backed voles, 19 woodland jumping mice) pooled across species. Among species there was variation in exploration time between individuals for each reproductive phase. Deer mice were consistently the most explorative with the exception of lactating females which were similar to lactating red-backed voles. red-backed voles consistently expressed the lowest amount of exploration time while woodland jumping mice were an intermediate between the two (Figure 3).

**Figure 1:**
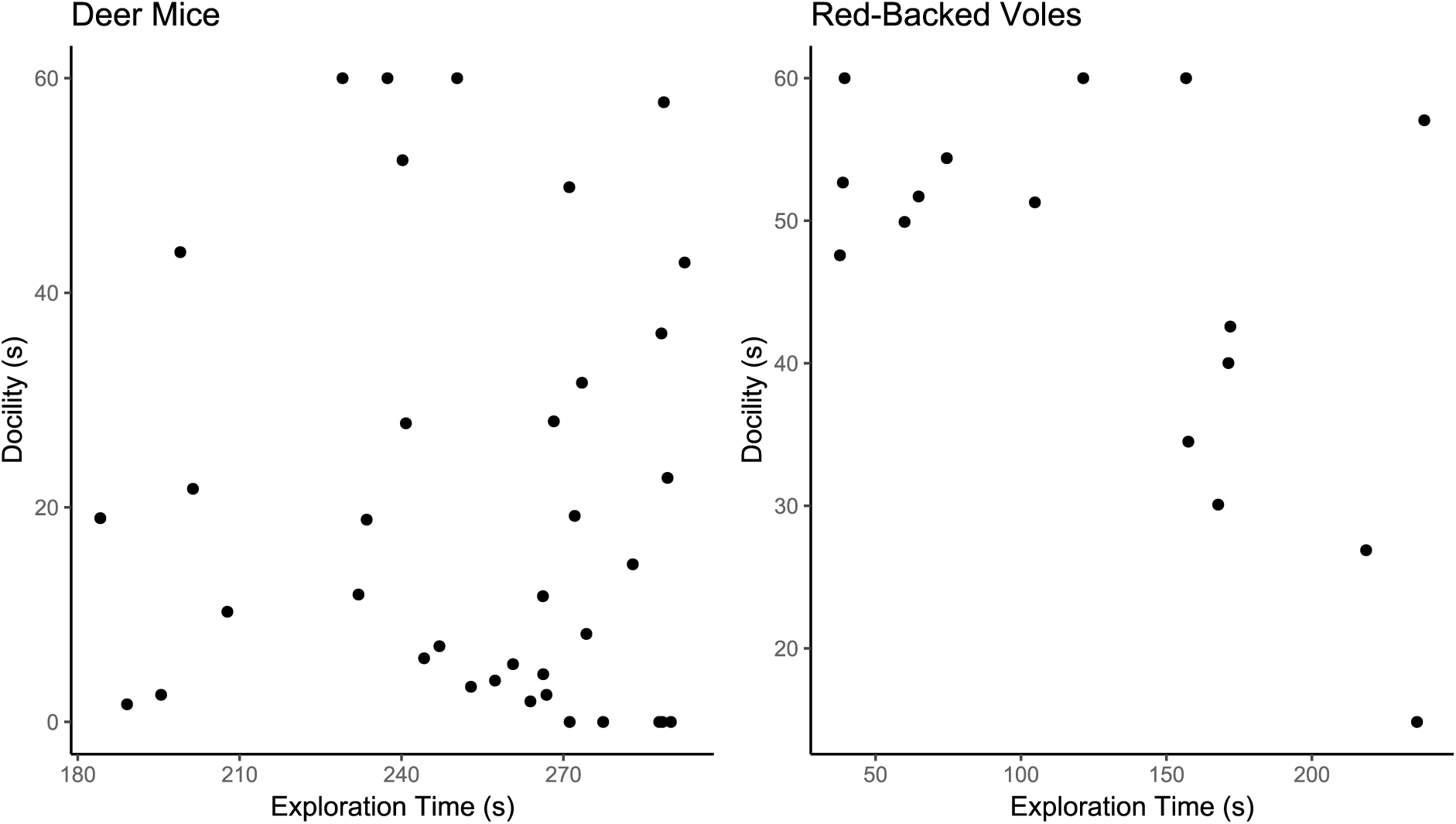
Scatter plots showing the relationship between total time spent expressing docile and exploratory behaviours in deer mice (r = -0.018, P = 0.26) and red-backed voles (r = -0.437, P = 0.091). Each data point represents the first Bag Test and Open Field Test an individual performed.

**Figure 2:**
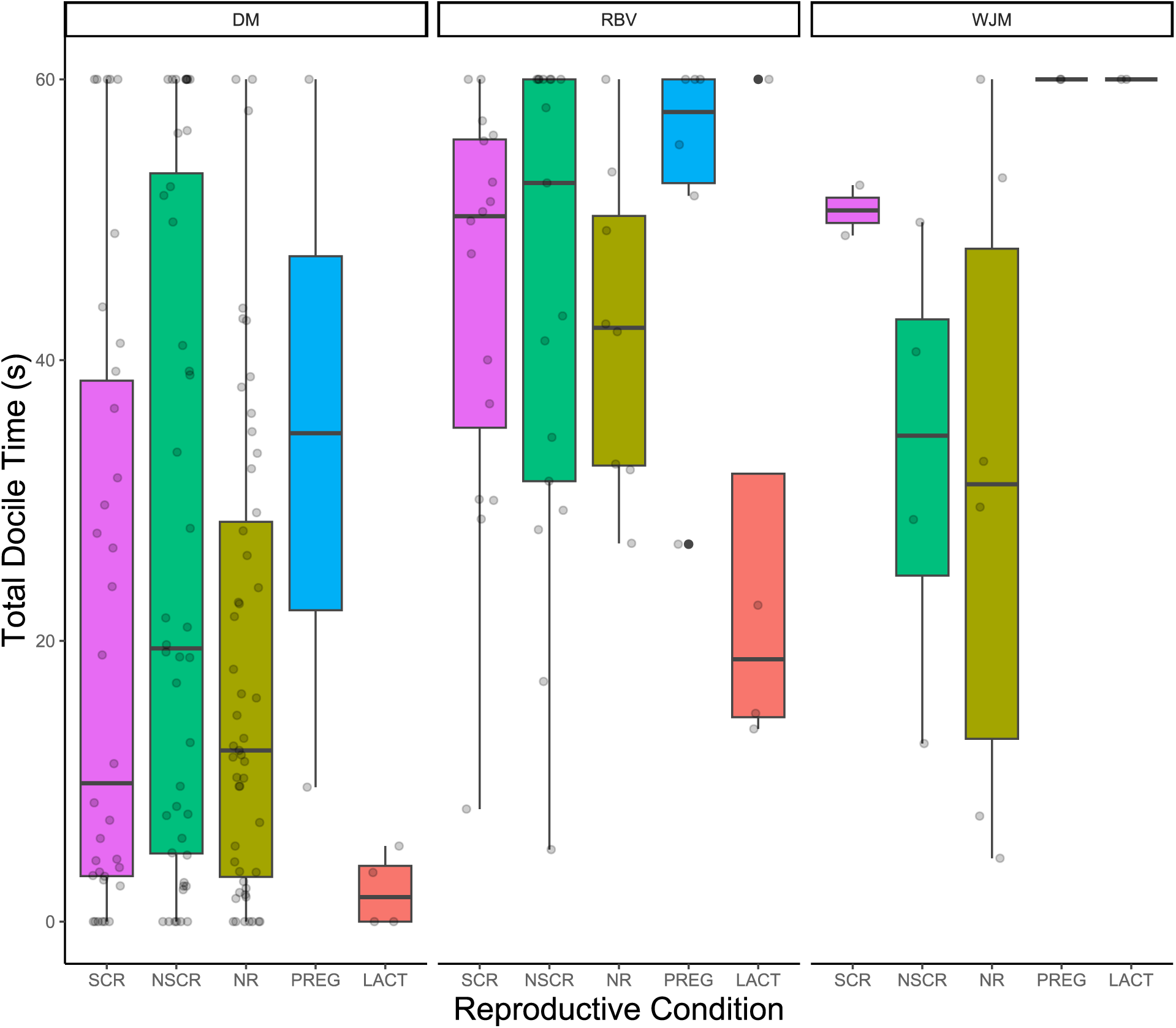
Boxplot showing the relationship between total docile time during the Bag Test comparing all three species, DM = deer mice, RBV = red-backed vole, WJM = woodland jumping mice, across all reproductive condition (SCR = Scrotal, NSCR = non-scrotal, NR = non-reproductive, PREG = pregnant, LACT = lactating) using individual ID as a random effect. Whiskers represent standard error; box lines represent the median time spent motionless across samples, boxes represent the range of motionless behaviour, and faded jitters represent the distribution of data.

**Figure 3:**
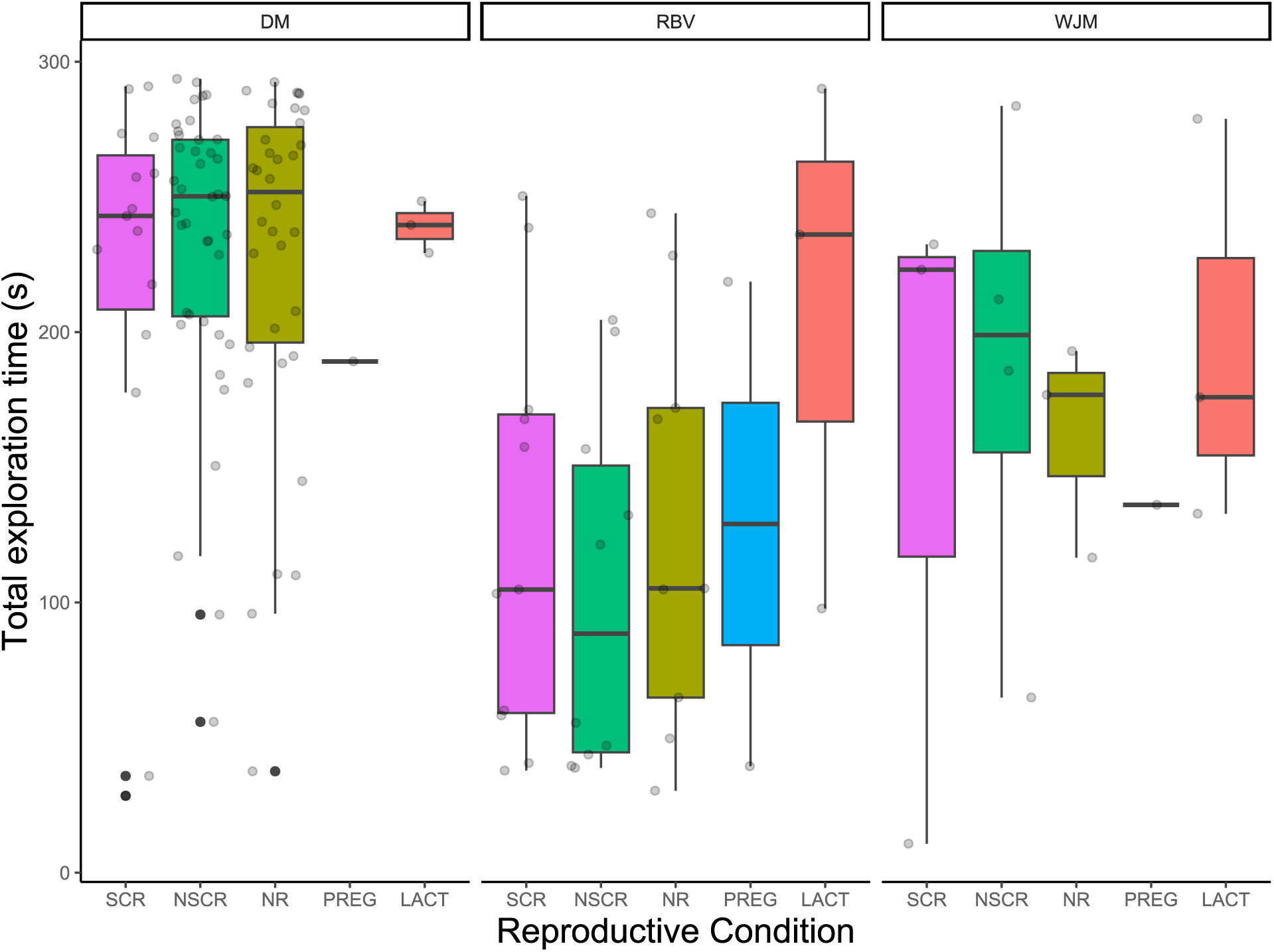
Box plots showing the relationship between exploratory behaviour (moving) during the Open Field Test for each species (DM = deer mice, RBV = red-backed vole, WJM = woodland jumping mice) for each reproductive phase (SCR = scrotal, NSCR = non-scrotal, PREG = Pregnant, LACT = Lactating, and NR = non-reproductive) using individual ID as a random effect. Whiskers represent standard error, box lines represent the median time spent expressing exploratory behaviours across samples, while boxes represent the range of time spent moving, and faded jitters represent the distribution of individuals.

## Discussion

We hypothesized that coexistence among sympatric rodents will promote divergence in behavioural strategy to accommodate resource competition, during the breeding season. We used differences in reproductive status between individuals of the same species to evaluate changes in behaviour throughout the breeding season. However, there was little effect of reproductive status on behaviour within species. Given potential differences in energetic costs at different stages of the breeding season (Gittleman and Thompson, 1988), we predicted measurable differences in docility and exploration between individuals of the same species. We also predicted that there should be a relationship between docility and exploration between individuals, such that more explorative individuals are less docile. However, we report no such significant relationship between personalities in our observed species.

We found significant differences in personality among species. Since docility and exploration are related to an individual’s tendency to disperse and engage with the environment, and thus, we predicted that more explorative individuals should be less docile (Martin and Réale, 2008; Réale et al., 2010). All three species observed within this study expressed a weak negative relationship between docility and exploration. We also showed differences in the position each species occupies along the predicted fast to slow continuum; deer mice were consistently more explorative and less docile across all reproductive stages than woodland jumping mice and red-backed voles. Meanwhile, woodland jumping mice were consistently more docile and less explorative than deer mice, with red-backed voles occupying relatively intermediate value between deer mice and woodland jumping mice, adhering to the predictions posited by the POLS hypothesis across species.

Empirical evidence to support the POLS hypothesis at the intraspecific level is mixed (Dammhahn et al., 2018; Royauté et al., 2018). Our findings further suggest that POLS between individuals of the same species may be more complex than the definition of the POLS currently allows (Royauté et al., 2018). We found little evidence that reproduction influenced exploration and docility between individuals in this study system. For males within all observed species, docility and exploration time remained consistent between scrotal and non-scrotal individuals.. In seasonally breeding rodents, changes in gonadal state are associated with energy use (Bergeron et al., 2011). Given the strong association between exploration and home range or foraging patterns (Gharnit et al., 2020; Spiegal et al., 2017) and the increase in hormones to illicit breeding, one would expect potential differences in behavioural strategies between individuals at different reproductive stages. Given the increased energetic demand for mate acquisition, males investing in reproduction through an enlarged gonadal state should have a higher level of exploration and a lower tendency to display docile behaviours (Hämäläinen et al., 2021). A low sample size for woodland jumping mice may explain some of the lack of differences we observed within this species. However, for deer mice, we propose two reasons for our results.

First, differences in movement behaviour may be negligible between breeding and non-breeding males. Although an altered gonadal state is associated with an increased energetic cost and altered hormonal state (Veitch et al., 2023), natal dispersal is necessary for juvenile and sub-adult mice (King, 1968; Rémy et al., 2011). Therefore, if potentially non-breeding individuals are dispersing at greater rates to alleviate competition, the trade-off in energetic investment between finding a mate and finding a new home may be comparable. Second, it remains possible that another factor strongly associated with dispersal and reproductive cost such as resource and habitat availability, is masking our results. Given the strong associations among mate acquisition, population abundance and resource availability (Bonte et al., 2012), it remains possible that there is a greater trade-off in the energetic costs of these population dynamics than the trade-offs presented solely by mate acquisition and reproductive investment during the breeding season.

We anticipated a decrease in exploration and increase in docility in pregnant and lactating females compared to non-reproductive females because of the costs associated with the risk of these behaviours. Indeed, several studies have observed an association between behaviour and reproductive status, including pheromone induced aggression (Martín-Sánchez et al., 2015), increased vigilance to protect against infanticide (Breedveld et al., 2019) and perhaps most significantly, hyporesponsiveness in lactating females (Chauke et al., 2011; Fleming and Luebke 1981; Lonstein 2005; Windle et al., 1997). Because we did not see the expected relationships between behaviours that may be anticipated with female rodents undergoing various stages of reproduction, it remains possible that some other factor is more strongly associated with these behaviours. For example, pregnant or lactating females may have an increased energetic cost associated with the production of milk or the development of young, but non-reproductive females may also be subject to selective pressures from potential predators or conspecifics that have a comparatively equal cost.

The POLS hypothesis also postulates differences in behavioural strategies and personality among species (Réale et al., 2010). Our results show differences in behaviour across observed species along the predicted continuum of deer mice – woodland jumping mice – red-backed voles. Competition among coexisting species may illicit variation in reproductive strategy and ultimately personality phenotypes. Despite apparent competition, there are few studies that aim to model personality in coexisting species (Wauters et al., 2019; Morris and Palmer, 2023).

Docile behaviours are often associated with behaviours that reflect risk taking behaviour, where a lack of aggression, increased predator avoidance, and a tendency to react to shock or stressful environmental stimulus remaining motionless may be considered docile (Boice et al., 1968; Martin and Réale 2008; Réale et al., 2000). Because of the association between docile personality types, and risk avoidance, docile behaviour is understood to influence seed predation and dispersal (Boone et al., 2022), pathogen spread (Zahdy et al., 2017), individual reproduction and survival (Goulet et al., 2016), and has an overall impact on how species interact with various environmental variables. We also saw a pattern across species in which less docile individuals, or those that expressed less time immobile, were more exploratory in the Open Field Test.

Exploration behaviour, often measured through an OFT is described as an individual’s tendency to move in a novel environment (Réale et al., 2007). The exploration-activity personality phenotype largely evaluates individuals that are more likely to engage in an action under novel stimulus, ultimately influencing dispersal and home range movement patterns (Réale et al., 2007; Boone et al., 2022).

Our results showing differences in behavioural phenotypes among species are consistent with the POLS hypothesis. Deer mice showed a lower level of docility and greater rate of exploration across all reproductive categories compared to red-backed voles and woodland jumping mice. Red-backed voles consistently expressed the greatest docility and lowest exploration, while woodland jumping mice expressed relatively intermediate values between deer mice and voles. While we anticipated red-backed voles should be more intermediate between deer mice and woodland jumping mice, these results are perhaps not surprising given these two species are closer along the fast-slow continuum, while the deer mouse has a much greater reproductive output and shorter lifespan compared to either. Given the differences in coexisting species that abide by the POLS hypothesis, we suggest that coexistence may be a driving factor in the development of alternative evolutionary strategies. Deer mice, red-backed voles and woodland jumping mice inhabit the same ecological niche, and population density is known to influence foraging and dispersal behaviour (Davidson and Morris, 2001). Given that each species is subject to the same environmental pressure, it remains likely that there is a benefit to alternative dispersal and movement strategies in sympatric species in the absence of niche partitioning of resources.

Animal personality can be influenced by an array of environmental conditions (Manning and Dawkins 2012). To help reduce the effects of environmental and handling variables, we used four preliminary factors associated with animal handling and capture including test start time, month of capture, cumulative capture count, and, for the Open Field Test only, if the individual received an ear tag before the test. Overall, we observed no significant correlation between start time and either docility or exploration. Similarly, we did not find any significant correlation between the month of capture and any of our personality traits. While seasonal variation in dispersal behaviours has been documented in laboratory settings using small rodents (Eccard and Herde 2013; Harrison et al., 2015), it is possible that the lack of seasonal variation we observed is the result of the timeframe of our study. Given that the majority of our samples occurred during the active breeding season it remains possible that more variation would be observed by comparing behaviours between spring and winter, or in months outside of the primary breeding season. It also remains possible that other seasonal environmental factors such as temperature or precipitation rate (Sergio 2003), or population dynamics including spatial competition or predator density (Bowler and Benton 2005) may instead have a greater impact on dispersal and movement behaviour, masking seasonal variation in our study.

Several studies have described a correlation between cumulative capture events and bold, or exploratory personalities (Boom et al., 2008; Boyer et al., 2010). Our results did suggest that the cumulative capture count did not have any significant impact on docility in any of our three species. However, we did find a weak positive correlation between cumulative capture count and exploration most prominently observed in deer mice, such that, individuals that were more likely to be captured, were more likely to be highly explorative in the Open Field Test, a commonly reported result (Johnstone et al., 2023; Boon et al., 2008; Boyer et al., 2010; Carter et al., 2012). Finally, we used a binary scale to compare recaptured individuals that already received an ear tag, to new individuals that receive an ear tag directly before entering the Open Field Test. Ear tagging, while common in live-capture research (Fokidis et al., 2006) will cause discomfort (Wever at al., 2017) and may potentially be a greater stressor than taking handling measurements. Further, individuals that receive an ear tag have a greater exposure to handlers prior to entering the Open Field Test, potentially further influencing the likelihood of expressed behaviours. Although there was a weak non-significant negative correlation in exploratory behaviour in deer mice that received a tag compared to those that did not, we did not see this trend for either red-backed voles or Jumping Mice. Despite the lack of statistical evidence, we suggest that increased handling time during live-recapture studies is likely to cause additional stress in an animal. Therefore, caution is advisable as small changes to behaviour may occur from prolonged handling time, or stressful measurements.

While we did not show evidence to support the POLS hypothesis between individuals of the same species, we show significant differences in behavioural phenotypes among coexisting species. Because species inhabiting the same ecological niche must engage in increased competition for resources, we further postulate that competition can regulate the expression of alternative behavioural phenotypes (Morris and Palmer, 2023; Sobral et al., 2023). Because personality may be a driving mechanism for environmental processes such as the distribution of seeds through fecal droppings (Réale et al., 2010; Morris and Palmer, 2023; Boone et al., 2022; Brehm and Mortelliti, 2022), we suggest that further evaluation into trade-offs in personality, population dynamics and changes in environmental structure will be valuable future research.

