## Supplemental data A for "Exploratory and risk-taking behaviours in coexisting rodents"

**Supplementary data A: Ethogram depicting common actions for all rodents in the 5-minute Open Field Test (OFT). Total exploration time, used to define the explorative personality phenotype in this study, is the summed total seconds of all actions involving locomotion, while time spent stopped or out of view from camera (obscured within the entrance) are considered non-explorative. Grooming behaviour, which also occurs in some individuals is considered a separate behaviour, and not included in the exploration personality phenotype.**

| Action | Definition |
| --- | --- |
| Total time spent walking, running, or jumping around. | Any form of forward locomotion |
| Time spent performing other locomotion (scratching, sniffing, scanning) | Individual remains stationary, with visible head movements or scratches the surface of the arena |
| Time spent stopped/doing nothing (visible) | Individual remains motionless in one location. |
| Time spent stopped (hiding) | Individual is obscured from the camera by hiding in the entrance |
| Grooming | Individual used hind legs to scratch themselves, licks paws, and/or uses front legs to scratch and comb face/body. |
