## Supplemental data B for "Exploratory and risk-taking behaviours in coexisting rodents"

**Supplementary data B: Ethogram for possible actions of all tested individuals during the one-minute Bag Test (BT). The docile personality phenotype consists of the summed total of seconds for all actions that involve remaining motionless (freezing/immobile behaviour). Non-docile actions are included as the individual attempting to flee or dig/scratch at the handling bag or other movement within the handling bag. For the purpose of this study, foraging and grooming behaviours are not considered along the docility personality phenotype.**

| Action | Definition |
| --- | --- |
| Flee, running, digging at bag | Individual attempts escape through bag by biting, scratching at bag |
| Freezing/immobile (visible) | Individual stops and remains motionless in one location |
| Grooming | Individual raises paws over head and continues to lick/scratch at self |
| Foraging | Individual continues to gather and eat seeds |
| Other locomotion | Individual is walking around bag with no distinct escape attempt |
